# Latent learning without map-like representation of space in navigating ants

**DOI:** 10.1101/2024.08.29.610243

**Authors:** Leo Clement, Sebastian Schwarz, Antoine Wystrach

## Abstract

Desert ants are excellent navigators. Each individual learns long foraging routes meandering between the trees and bushes in their natural habitat. It is well-known how the insect brain memorizes and recognizes views, and how this recognition can guide their way. However, little is known about the rule that guide spatial learning in the first place. Here we recorded the paths of desert ants navigating in their natural habitat under various displacement conditions. We demonstrate that ants learn *continuously* the routes they travel and memorize them in one trial, without the need for reward or punishment, and even if these routes are meandering and do not lead to places of interest: a concept called ‘latent learning’, which is typically associated with the formation of map-like representation in vertebrates. Yet, the failure of ants to solve simple artificial navigation tasks -even with the goal being clearly visible- reveals that they relied on egocentric visual memories without map-like representation of the surrounding space. Our results unveil the rules governing the formation and recall of latent memories. A model shows that it can be implemented in the insect’s Mushroom bodies brain area through dynamic interactions between short- and long-lasting memories.

## Introduction

Many animals must acquire spatial knowledge to navigate the world. But spatial learning is a peculiar form of learning. In some instances, it fits the more general learning theory, where learning depends on rewards or punishments (i.e., an unconditioned stimulus US). For instances, animals, from maggots to mammals can associate a specific location (the conditioned stimulus CS) with food^1–3^ (US reward) or an aversive event ^4–6^ (US punishment). However, such discrete rewarding or punishing events only occur sporadically along the journey of a navigator. Most of the time, navigation is a reward-less continuous task. Nonetheless, animals can acquire spatial knowledge over large region of space without apparent reinforcement ^7^, a feat so-called latent learning ^8–10^.

Latent learning has been mostly studied in vertebrates and is typically assumed to involve – or be associated with – the continuous reconstruction of a map-like representation of space ^11^. To be built, a map-like representation requires the interaction between path integration and the learning of visual scenes ^12–15^. As for vertebrates, insect navigators such as ants and bees are known to rely on both path integration (PI), and the learning of visual scenes ^16–24^. But whether they perform latent learning while navigating is unclear, and whether they form map-like representations is much debated ^3,25–29^. Here, we investigated those questions through a series of manipulations with desert ants (*Cataglyphis velox*) navigating in their natural habitat. Our results demonstrate that ants learn the routes they travel in a continuous fashion, and without the need of reward or punishment, that is, they acquire spatial knowledge continuously through a latent learning process. However, we show that these memories are not used to form a map-like representation of the space around them. Instead, latent memories are used to define moment-to-moment egocentric decision, and ants use a dynamic gating mechanism to ensure the appropriate storage and recall of such continuously building memories. We show how such a continuous mechanism for latent learning can be implemented in the neural circuit of the insects’ mushroom bodies.

## Results

### One trial is sufficient to learn a route

We first investigated whether ants would memorize a route encountered only once in a distant unfamiliar test field. To do so, ants trained to a 7.0 m familiar route were captured upon departure from their usual feeder, that is, just before starting their homing journey, and released on a distant unfamiliar test field (Fig. 1A, S1B). These foragers are called full vector (FV) ants as they possess a path integrator (PI) homing vector, pointing to the feeder-to-nest compass direction. Unsurprisingly, when released at the centre of the test field, these FV ants followed their PI vector direction (Rayleigh test with PI as theoretical direction p: FV< 0.001; Fig.1 B,C, third row). Upon reaching the rim of the test field (3.0 m) the ants were captured again and released directly inside their nest (N) (Figure 1 A, B first column). On their subsequent exit of the nest, the same individuals were free to run to the feeder and back along their familiar route. This time, those foragers were caught just before entering their nest and released for the second time on the distant test field. These ants are now in a so-called zero-vector state (ZV) because they have already run out their PI homing vector, and must rely on visual memories to guide their path ^30,31^. Remarkably, when released on the test field –which they have experienced only once – these ZV ants followed the same direction as during their first trial as FV ants (Rayleigh test with PI as theoretical direction p: FV< 0.01; Fig.1C, first column). This shows that they relied on memories built during the first trial. To check whether these ants had learned a route rather than just a general direction of travel, we compared the shape of their paths displayed on the test field using dynamic time warping analysis (DTW). Between the first training trial (FV) and the subsequent test trial (ZV), paths shapes were more similar within individuals than between individuals (Figure 1D, W = 11, p < 0.05). This shows that one trial is sufficient for ants to learn an idiosyncratic route in a novel environment (Figure S2, first row).

**Figure 1.**
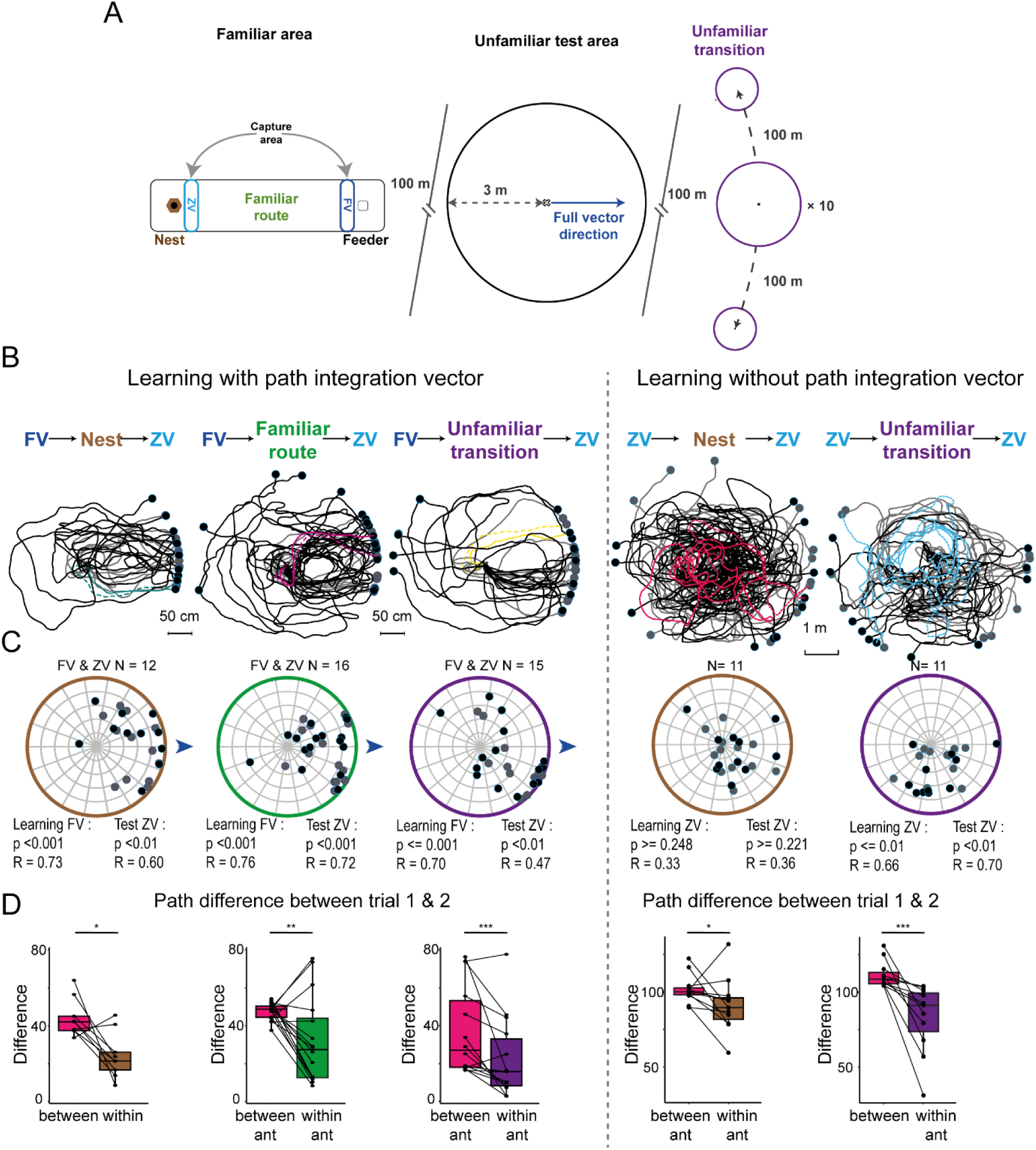
Ants learn and recall their route even if it leads to nowhere. (A)Scheme of the experimental set up. Familiar route consisting of a straight route of 7.0 m enclosed in a corridor (7×1.5 m). A feeder is placed 7.0 m away from the nest at least 24h before the experiment. (Unfamiliar test field 100 m away from the familiar area. Unfamiliar transition areas placed at least 100 m away from the test field and 200 m from the closet nest. Trajectories of *C. velox* ants recorded on the test field. For their first trials ants were released as full-vector (left panel) or zero-vector (right panel) at the centre of the test field. Ants were then released either in their nest (brown), on their familiar route (green) or on another unfamiliar terrain (purple) and captured again as ZV ants. For their second trial, these ZV ants were released again at the centre of the test field. (B) shows the training path (grey) and test path (black). Within each condition, an example-coloured path has been emphasized for the training (dashed) and test (solid) path. The colour of the path corresponds to the ant identity. (C) is a circular plot showing the average circular vector calculated over a portion of the path during the first trial (grey) and the test (black). Each dot is the mean direction taken and the mean resultant vector (i.e., a point closer to the periphery indicates straighter paths). (D)shows the results of the similarity analysis (DTW). This analysis was made between the first and second trial (see methods). Each ant’s path has been cut into segments (0.9 m for A path and 2.4 m for C). Each segment has been compared with the closest neighbour segment of the subsequent trial to estimate their similarity. We did this analysis by comparing two trials of same ants (within ants, right boxplot) or between ants (left boxplot).

### Learning a route needs no reward

In the previous experiment, ants were released inside their nest right after their first trial on the test field. We hypothesized that entering the nest may constitute a reward for the ants to consolidate memories of the route they just had experienced before entering the nest. To test for this, we replicated the experiment but, this time, ants were prevented to enter their nest. After their first trial as FV ants on the test field, foragers were released either on their familiar route (Figure 1B, second column, F) or in another distant unfamiliar location (Figure 1B, third column, U). In both cases, the ants were free to run off their PI vector before being captured again (as ZV ants) and released on the test field for the second trial. Here again, ZV ants were well oriented, and their paths resembled mostly their own previous FV paths (Figure 1B, C, p< 0.01), demonstrating that they had learned an idiosyncratic route. Thus, entering the nest is not necessary for ant to memorise a route. Ants can learn a route in a single trial and subsequently recall it regardless of whether they have experienced their nest, a familiar route or another unfamiliar terrain in between (Figure S2). Note that the latter appears somewhat maladaptive, as ants are keen to recapitulate their freshly learnt route even though this route had led them to nowhere but another unfamiliar location (but see discussion).

### Learning a route needs no path integration vector

We next wondered whether route-learning occurred because ants were running down their PI home vector (feeder-to-nest compass direction) during the first trial on the test site. Indeed, running down the home vector indicates to the ant that she is moving toward the nest direction, and thus might act as a reward signal for learning the current route ^32^.

We captured homing ants on their familiar route just before they reached their nest, that is, as ZV (and thus deprived of PI homing vector) and released them on the test field for their first trial (Fig. 1A,). As expected, these ZV ants displayed a typical search path ^33–36^ on the test field (Fig. 1B,C, last two columns, Fig.S3A,B first column). They meandered in all directions, walking in average 21.5 m (sd = 9.5 m) before reaching the rim of the circular test field (3.0 m radius). Once the foragers had reached the rim, they were captured and subjected to one of the two following experimental groups: either they were released inside their nest (Fig. 1B, C, 4th column, N) or released for ∼30s in another unfamiliar terrain (Fig. 1B, C last column, U) before being captured again. Once released on the test field for their second trial, the ants did something strange: they tried to follow their previous meandering path as if it was a route: both groups showed a strong idiosyncrasy by reproducing characteristic signatures of their own previous meandering path (ps < 0.05, Fig. 1D, last two columns). Consequently, even without running down their PI vector, ants had memorized their previous meandering search and used this memory to guide their subsequent trip, even though this previous trip consisted of a meandering search in an open test field!

Overall, we are left with the conclusion that ants navigating in unfamiliar areas continuously and rigidly learn the route they are travelling and subsequently recognize and recall these memories to guide their path, even in the absence of any sort of external or functional reward, and independently of the dictates of their path integrator.

### Compulsive route following rather than shortcutting

The fact that ants followed their route even though this route meandered into circles within a small open field (Figure 1D, S3) appears somewhat puzzling for its apparent pointlessness. We tested whether ZV ants would optimise their path over trials by releasing them up to ten consecutive times at the centre of the test field. Each time the homing ants reached the edge of the 3.0 m radius open field, it was transferred back to its nest. From a spatial point of view this task is dead simple. It requires to walk towards the edge of the test field to reach the nest; just moving straight in any direction for 3.0m would suffice. Nonetheless, the ants showed no improvement over trials (Figure 2A, D, S2, S3), they carried on following a meandering path, often pointlessly repeating the same looping segment again and again (Fig 2, S3). All ants tested had a cookie crumb and there is no doubt that they were motivated to reach their nest. Nonetheless, they did not learn to shortcut towards such a goal even though the rim of the test field was in plain sight. What’s more, ZV ants trained on a similar regime (also for 10 trials), but systematically released at other unfamiliar locations (instead of the nest) between trials also showed consistent route following on the test field with no improvement (Figure 2, S3), even though these routes literally lead to nowhere familiar. We are left with the conclusion that ants navigation is *not* based on computations grounded on a reconstruction of the surrounding space, but on compulsively following ever building egocentric route memories.

**Figure 2:**
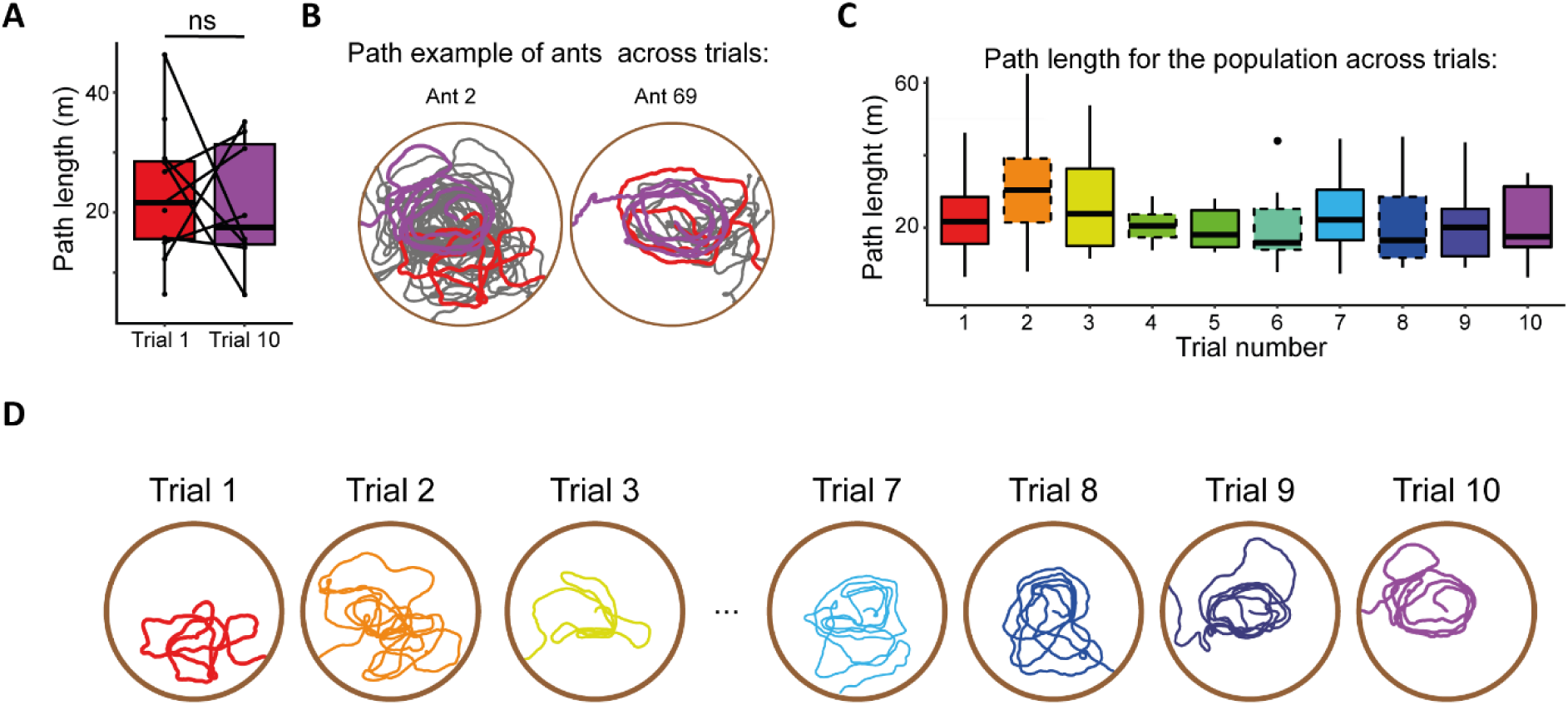
Ants do not optimize the distance covered trials after trials. Homing ZV ants are released on the test field for 10 consecutive trials, and replaced inside their nest as soon as they reach the rim of the test field (3m radius). (A) The distances covered across trials one and ten are similar (p > 0.05) (B) Example paths of two individual highlighting that the distance covered has not improved between trials one (red) and ten (purple). (C) Distributions of individuals’ path length across trials. (D) Example paths of successive trials of a same individual, highlighting the repetition of a same looping trajectory across trials (see supplemental material for more examples).

### Ants recall memories of the previous, but not the current, trial

We reasoned that if learning is continuous, and the memories formed are egocentric route instructions, it is crucial for the ant to prevent the instantaneous recall of these ever-building memories. Otherwise, the ant would not be able to distinguish what is familiar and what is not, and would aimlessly follow a self-building route as she walks.

Our result show indeed that ants learn novel memories but don’t recall them during trial 1. This is well illustrated by the search patterns displayed by ZV ants on the test field. The first and second trial on the test field have very different characteristics. During their first trials, ants displayed a typical systematic search: exploring novel areas as the search unfolded (Figure 2A, B, first column, third row; grey path). Contrastingly, on their second trials, ants tended to explore less areas (Fig. 3A, B first column; exploration index between path 1 and 2: p-value: Nest < 0.01, Unfamiliar > 0.05) and walked along portions of the route they had already travelled, sometimes getting stuck into walking along a same looping trajectory multiple times in a row (Fig. 3A, B black paths, Fig. S3A, B trial 1 and 2). The paths during the second trials shows more self-overlapping portions than the first trial did (Fig. 3 A, B second column; self-difference index: DWT N = 11, T = 3, ps < 0.05). In other words, during the first trials, ants were performing a systematic search (as they typically do in unfamiliar areas); and during the second trials, ants were following the route they had memorised in the first trial (Fig. 3A, B black paths, Figure S3A, B). This confirms that the route memories acquired during the first trial were learnt, but not recalled during the first trial. What then, triggered the recall of the route memories?

**Figure 3.**
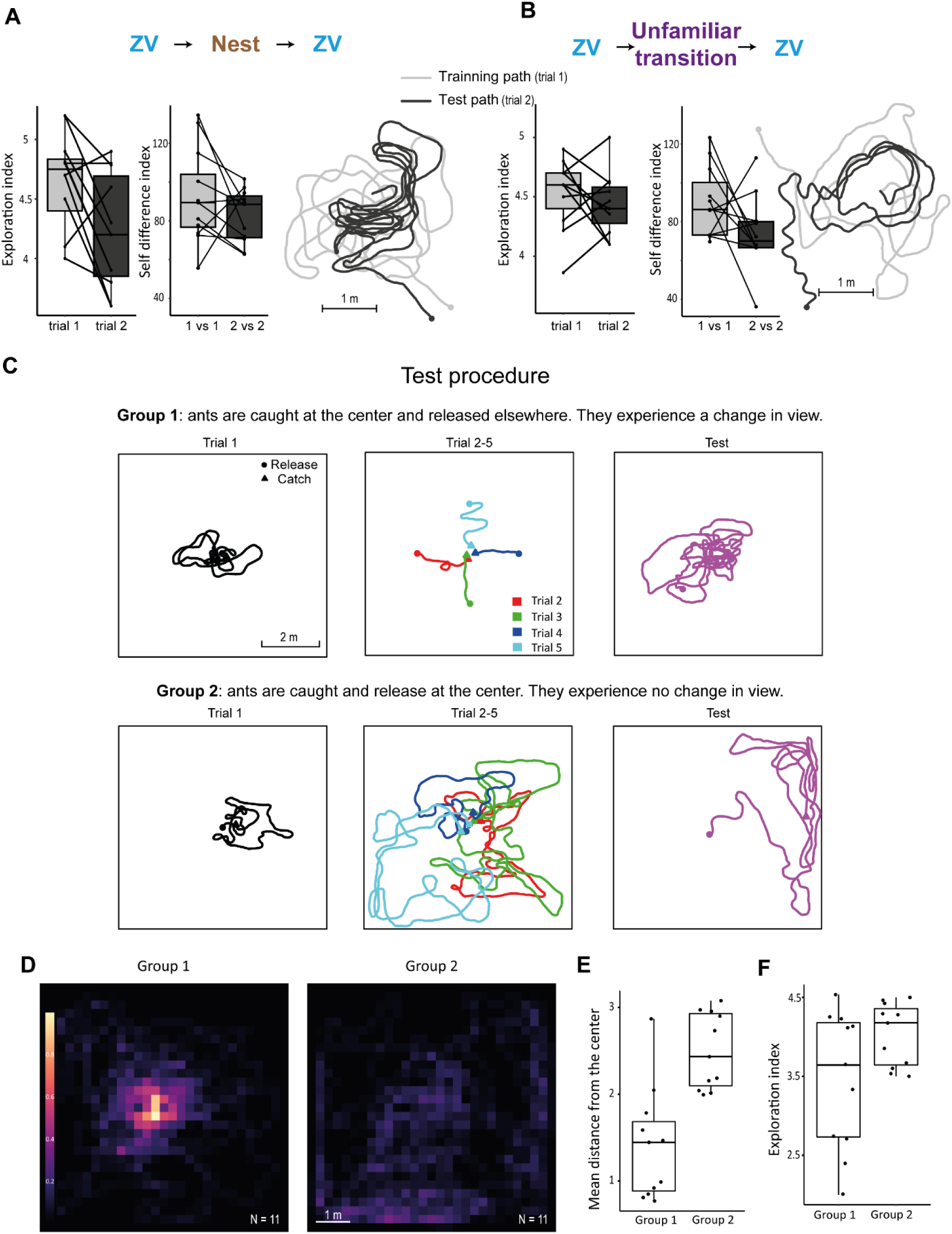
A discrete change in view triggers memory recall. (A, B) Path analysis for both visual transitions conditions. Exploration index: the first 10 m of the path are discretized into an X/Y matrix with a resolution of 25 cm. The exploration index is then calculated by dividing the number of explored cells divided by the length of the path. Self-difference index indicates how much ants tended to repeat signatures of their current path. Each ant’s path has been cut into segments of 2.4 m and each segment has been compared with the closest neighbour segment to estimate their similarity (DTW, see method). For each ant, we used the median score of all its route segments. Example paths showing that ants tend to explore during trial 1 (grey) but repeat their route segment guided by visual memory during the subsequent trial (black). (C) Example paths illustrating the test procedure. For the first trial, ZV ants are released and captured at the centre of the test field. For the next four trials, ants were captured at the centre, and either systematically released at a different location (group 1) or at the centre again (group 2). During tests (the sixth trial) ants are recorded without interference. (D) Density heat map at the population scale of the ants’ position during the test for group 1 and 2. The density maps have been done on the first 28 m walked of each ant. More examples of paths are presented in Figure S3C. (E) Distribution of the mean distance from the centre reached during the test. (F) Distribution of exploration index, the first 28 m of the path have been discretized into an X/Y matrix with a resolution of 25 cm. The exploration index is then calculated by dividing the number of explored cells divided by the length of the path.

### Recall is triggered when ants experience a discrete change of viewpoint

In our experimental manipulation, something happened between trial 1 and 2 that led to the recall of the memories acquired during trial 1. It could be the fact that the ants were released in another location (either their nest or another unfamiliar terrain) and thus experienced a discrete change of view in between both trials (hypothesis 1) or the fact that ants spent time in the catching vial (∼60s) between both trials (hypothesis 2). To test for these two non-mutually exclusive hypotheses, we performed the following experiment. We captured a new cohort of ants as they exited their nest (ZV ants), gave them a cookie to trigger homing motivation and released them at the centre of the test field for the first time. As expected, these ants displayed a tight systematic search at the centre of the test field (Fig. 3 C trial 1, S3 A,B). If, during their exploration, ants would venture further away than 1.8 m and then return to the centre of the test field, they were captured again (at the centre) and kept for 60s in the vial. From there we created two experimental groups: some ants were released at a different location in the test field and thus experienced a discrete change of viewpoint (group 1), while the others were released again in the centre of the test field, that is, where they had just been caught, and thus spent time in the vial but experienced no discrete change in view point (group 2). As in trial 1, ants of both groups were free to explore and were captured again if reaching the centre of the test field. After sixth trials of this regime, ants were recorded while exploring the test field without interference.

Remarkably, ants of group 1, who were released at a different location and thus had experienced discrete changes of views between catch and release, concentrated their path densely at the centre of the test field (Figure 3E; mean distance from the centre: group1 vs group2 one-tail test = p < 0.001). This is what is expected if ants are recalling their memories of the previous trials, which were concentrated at the centre of the test field from the first trial onwards. Remarkably, this strong attraction to the centre is visible from trial 2 onwards (Figure 3C, trial 2-5) and confirms that ants are recalling memories rather than pursuing a systematic search (Figure 3D, F; exploration index: group1 vs group2 one-tail test = p < 0.05). Inversely, ants from group 2, who were systematically caught and released at the centre of the test field, and thus did not experience a discreet change of view, searched mostly around the peripheries of the test field (Fig. 3D, E). This is what is expected if ants are not recalling their memories of the previous trials but instead simply pursuing their systematic search, which is indeed expected to expand toward the periphery with time^33,34^.

Together, this demonstrates that spending time in a vial does not trigger the recall of the previously formed route memories; however, experiencing a discrete change of location does. Perhaps counter-intuitively, this makes functional sense. By doing so, the ants are safeguarded against the risk of recognizing a currently experienced novel viewpoint as familiar when it is not. Instead, novel routes experiences will always be treated as unfamiliar while being explored since no discrete change of views occurs, enabling other mechanisms, that are adapted to unfamiliar environments (such as path integration or a systematic search) to control guidance. A discrete change of view ensures that the ant is no longer perceiving the previously learnt route (or route section), thus the inhibition of the recall of these memories can be safely lifted.

### Neural implementation of latent memories and recall

We investigated how the observed dynamic of learning and recall could be implemented in the insects’ neural circuit using a modelling approach. Route memories, which are mainly visual in solitary foraging ant species ^28,37–43^ are stored in a brain area called the mushroom bodies (MB) ^44–46^. Each perceived view triggers the sparse activation of a unique set of MB Kenyon’s cells (KC) and learning occurs through the modulation of the output synapses of these KC’s to MB output neurons (MBON) ^46,47^. As a result, MBON activity provides information about the familiarity of the current view. Insects possess multiple MBONs mediating parallel memory banks with different proprieties ^48,49^, such as: short vs. long-lasting memories ^50–53;^ ‘views of the way out’ vs. ‘views for homing’ ^46,54^, or in the context of olfaction, odours associated with sugar vs. odours associated with punishment ^55–59^. The recall of memories is then usually controlled by a complex gating system of local (between MBONs) and external modulations of the MBONs, typically conveying information of the animal’s current state and needs ^46,55,60–62^.

Based on this type of connectivity, we present here a simple circuit that explains the dynamic of visual learning and recalling in ants, while strictly respecting known features of the interactions between different MBONs. Our model assumes that:

- A long-lasting ‘persistent memory’ (PM) and a short-lasting ‘transient memory’ (TM) of the route experienced are continuously built in parallels in different MB lobes compartments and thus conveyed by different MBONs (as observed in ^50–53)^.
- The MBON conveying the PM controls steering behaviour (as in^47,54,63^), and the MBON conveying the TM inhibits the expression of this control through direct lateral inhibition (as in ^55,64^) onto the PM MBON (Fig. 4A).
- The TM memory is reset if the ant enters its nest, or if a discrete change of location with a novel view is experienced. The reset can occur by having the input synapses of the TM MBON switched to default again (as in ^48,60^). Note that information about such a ‘novel discrete change of view’ could be obtained by having the TM MBON activity itself providing (direct or indirect) modulatory feedback on its own input synapses (indeed, because learning is continuous, strong unfamiliarity can only arise sporadically given a discrete change of view that hasn’t been experienced during the current trial). More likely, such information could be implemented as a third party MBON, acting as ‘novelty detectors’ that send a lateral connection to modulate the TM MBON input synapses (as in ^65^).

**Figure 4.**
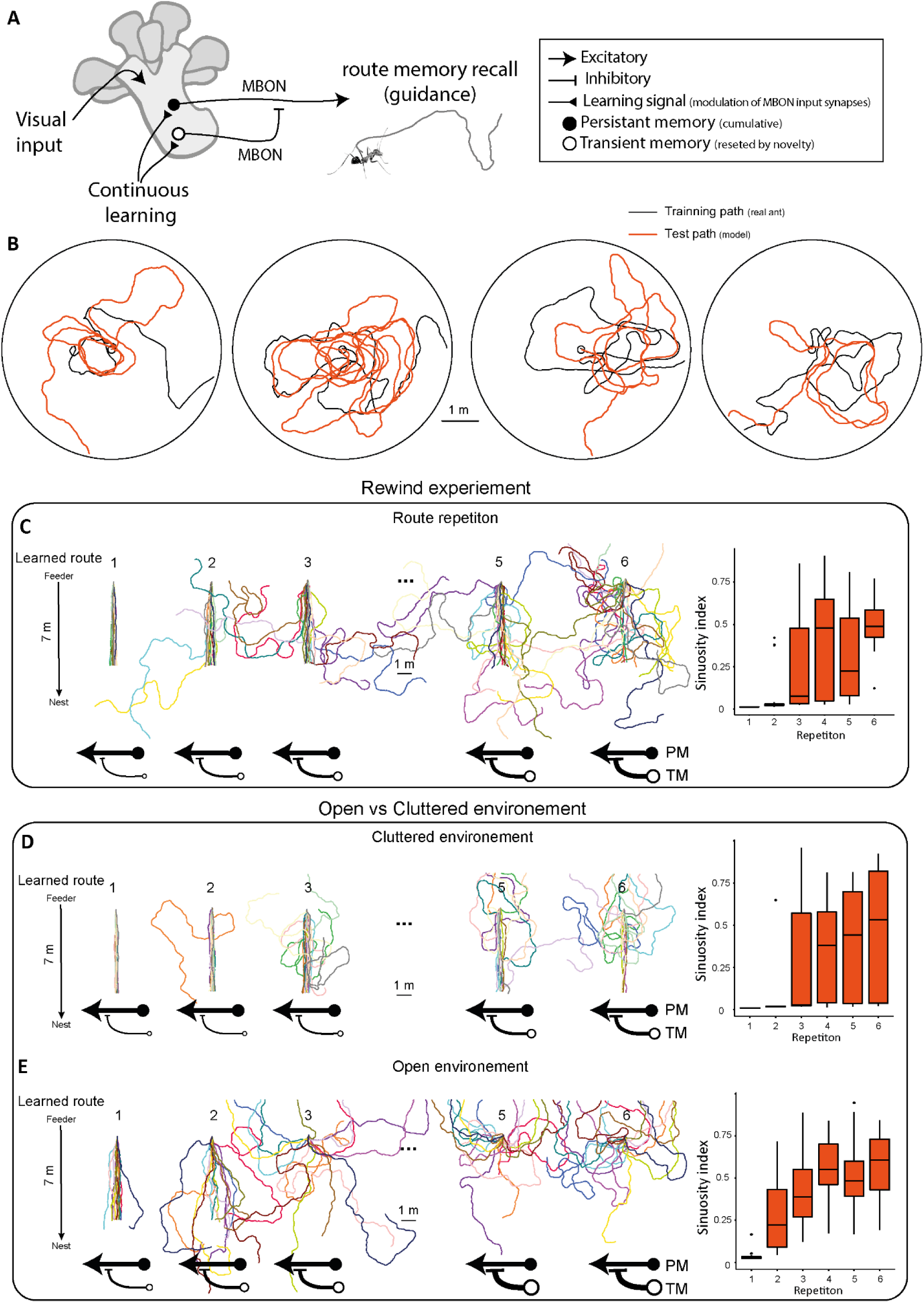
A simple circuit captures the dynamic of route memories in ants. (A) Scheme of the model. Model with a continuous build of two memory banks (persistent long-lasting memory (PM) and a transient short-lasting memory (TM) that inhibit each other. (B) Example paths of test (trail 2, orange). The model has been trained once on real data (black path) and then we simulated 10 replicas of the test trails (orange) guided by memory bank acquired during training, we selected parameters to prevent the model to being more efficient in route following than our real ants (Figure S4). (C-E) Reproduction of the experiment made by Wystrach et al., 2020^70^. First 20 ant agents are trained 50 times to establish a route between the feeder and the nest located 7.0 m away. During training, ants navigate with a home vector and progressively update their memory banks. Under each path, the thickness of the black arrow corresponds to the relative strength of the TM. While the thickness of flat head arrow is relative to the strength of the TM. (C) Test the effect of repetition of route on route following. After 6 repletion the path meander away significantly (sinuosity index trial 1 vs 6, N= 20: V = 210, p<0.001). (D, E) Test the effect of the structure of the panorama: cluttered (D) and open (E). (C-E) Boxplot shows the distribution of sinuosity according to the repletion.

As a proof of concept, we implemented this circuit in a navigating agent. The agent is guided by 3 forces (on which noise is added): 1-path integration, that tends to make it walk towards the ZV point; 2-Inertia, that tends to make it carry on straight (on which we add noise); 3-current visual familiarity, as given by the output activity of the PM MBON (resulting from the circuit described above, Figure 3A) for the current location and heading (because visual recognition in ants is rotationally dependent ^66^). This force tends to make the agent turn in the most familiar direction at this location (as shown in ^63^) and thus follow its familiar route. The driving force of the current visual familiarity is weighted as a function of the PM MBON activity itself ^67,68^, so that a strong familiarity yields the agent to pursue almost perfectly along its route direction, while a weak familiarity lets path integration and noise more control on the behaviour.

Despite continuous learning, this model explains the absence of memory recall while exploring a novel environment. This is because during the first trial in a novel environment, both TM and PM are building memories in parallel (Figure 4A). Any views that have already been experienced (for instance when performing a loop) is simultaneously ‘recognized’ by both memories, and thus triggers the equivalent activity in both PM and TM pathways. The signal of the TM MBON matches and thus suppresses the activity of the PM MBON, which therefore does not influence behaviour. In other words, the route memories of the current trial are not recalled and do not impact the behaviour, letting instead path integration and noise having full control over the agent’s movements: a search pattern naturally emerge. During the second trial, however, the transition (in the nest or in unfamiliar terrain for real ants) resets the TM, which therefore, no longer inhibits the visual familiarity signals of the PM, enabling the recall of the previous trial route memories when the familiar views are encountered. As a result, when the agent during trial 2 is on a portion of its route effected during trial 1, views are recognized, and the non-inhibited PM pathway leads the agent to carry on and recapitulate its previous trajectory (Fig 4A, S4A).

Interestingly, this simple model captures further puzzling results from the current and previous works in ants, even-though it was not designed for it. First, it explains why ants during their trial 2 repeat portions of their first trials multiple times but not indefinitely, ensuring that the agent (like the ant) will eventually ignore their maladaptive route and reach the border of the test field (Figure 4B, S4A, C). This is because, as the agent navigates its second trial, the TM is building up again and eventually catches up with the PM if a route segment is being repeated. In other words, as the agent repeats a looping route portion, the inhibition of TM on PM yields it to increasingly ignore the repeated segments and therefore free himself from running into circle. This property turns out to be vital to prevent the agent getting stuck in an infinite loop and captures the wide variation observed across individuals in our experiment (Figure 4B, S4A). Second, the model captures the evidence from previous work that ants forced to repeat a route segment several times consecutively eventually stop following the route and increasingly meander away (Figure 4C)^69,70^. Indeed, as the agent repeats its route segment, the TM memories build up and increasingly inhibit the PM (Figure 4C). Third, and perhaps most impressive, our model captures the fact that this disruption of route-following through repetition occurs more quickly in a visually open environment than in a cluttered environment (as shown with added proximal objects on the side of the route)^6^ (Figure 4D, E). This is because views change slower with displacement in an open environment, and therefore, this longer lasting exposition to the same views (repeated activation of the same KCs) leads the TM memory to build up and thus inhibit the PM memory more quickly than in cluttered environments (Figure 4D, E).

## Discussion

We demonstrated that *Cataglyphis velox* ants navigating in their natural habitat memorize a route through a novel environment in a single trial without reward and independently of their path integration state. Foragers subsequently recall these route memories regardless of whether they have experienced the nest, a familiar or unfamiliar terrain in between. In other words, these navigating insect learn all the time. Such continuous learning, without any form of external or functional reward, echoes the so-called ‘latent learning’ observed in vertebrates spatial learning ^8–10^. Indeed, as for vertebrate, learning is here truly ‘latent’ in the sense that it is continuous but not used immediately. Also, it demonstrates that the smaller brain size of insects too can store a continuous flow information as it is experienced, and without saturating (see ^71^, for an example of this can be implemented in neurons).

In the vertebrate literature, latent learning is associated with the reconstruction of map-like representation of space ^11^. However, our results highlight a profound difference of the representation of space between ants and vertebrates. Whereas vertebrates spontaneously optimize their paths between a start and a goal location – a signature of map-like representation ^72–75–^, ants did not, instead, they did the contrary (Figure 2, S2A). ZV ants released – up to ten consecutive times – at the centre of a 3.0 m radius open field with ample visual information did not learn to simply walk to the periphery, where they would systematically be transferred to their nest (Figure 2, 3A, S3). Instead, they follow their route memories even when it led them to walk ‘uselessly’ into circles on the test field (Figure 2,3A, B, S3; see also ^76,77^), with no improvement over trials (Figure 2, S3B) and despite the fact that the goal was in plain sight. This aberrant, non-functional behaviour seems impossible to conciliate with the idea that ants build a unified representation of the space around them. Instead, it reveals the heuristics that are at play: homing ants continuously learn to associate the egocentric scenery perceived to the motor command performed, even if the movements effected leads to nowhere. In natural conditions, this heuristic is perfectly functional as path integration guide the early trajectories of the naïve insect in a functional way, thus acting as a scaffold for learning routes that lead effectively to goals such as the nest or food sources ^38,78^.

Finally, our growing knowledge of insect neural circuits enables us to explains how a wide variety of the behaviours observed can emerge without building such a unified representation of space^46,47,47,63,79–82^. In the same vein, we showed that the peculiar dynamics of latent learning and recall observed here can result from a simple interaction between the expression of two memory banks in the Mushroom bodies (Figure 4). The different temporal dynamics of these memory banks ensure that route memories are not recalled inappropriately, and most importantly, safeguard the insect of being stuck walking along a loop infinitely, a risk that exists *only* if using egocentric route-following heuristics without map-like representation.

Whether latent learning has evolved for navigational reason, or whether it is equally present in other context of insect learning involving interaction across the MB lobes ^53,55,64^; whether it is ancestral to insects and vertebrates common ancestor, or whether it has evolved independently in species that needed great spatial skills, remains to be seen.

## Methodology

### Study Animal & Experimental site

All experiments took place within an open semi-arid landscape near Seville (Spain) from Sep. to Oct. 2021. We had the opportunity to work with the thermophilic ant *Cataglyphis velox.* Foragers, rather than using pheromone, are known to navigate solitarily relying mainly on learnt terrestrial visual cues and path integration (PI). For the main experiments, four nests have been located and used within an area of about 200 square meters. Each nest has been enclosed in a corridor (7.0 × 1.5 m), with a feeder placed 7.0 m away from the nest and foragers were free to familiarize themselves with the route for 24h before being tested.

### Experimental procedure and data collection

We investigated whether ants would learn a novel route in a distant unfamiliar test field (Figure 1A, S1B) during their homing path. For all experiments, the test area was covered with a gridded string pattern (1×1.0 m) to record the path of the ants.

#### First trial as Full vector ants

Ants were captured at the beginning of their homing path along their familiar route corridor. Hence, those foragers are called full vector ants as they possess a homing vector (FV) pointing to the food-to-nest compass direction. Ants were trained in an unfamiliar test field (TF) at least 100 m away from their usual route (Fig. S1). Paths were recorded until the foragers reached the outer rim of a 3.0 m radius circle. When they reached the rim, ants were captured and released in one of three distinct visual transitions: (1) directly inside their nest (N); (2) along the ant’s familiar route (F) therefore in presence of familiar terrestrial cues; (3) or in an unfamiliar location (U, Fig.S1B) 100 m away from the test field (Figure 1A). Crucially, for the subsequent trips in the TF ants were tested without homing vector (ZV ants). Hence, they could not rely on their PI vector but only on visual memory acquired during the previous trail. To do so, we let the ants run off their vector (approximately 3.0 m) either within the unfamiliar (U) transition or along their familiar route (F). For the group of ants released inside their nest (N), we observed the nest entrance, followed foragers until they picked up a food item at the feeder and finished their inbound trip and captured them just before the nest entrance. Hence, they were tested at ZV ants.

#### First trial as Zero vector ants

To test whether ants would learn a route on the unfamiliar test field without PI homing vector, homing ants were caught on their familiar corridor just before entering their nest and thus no longer possess a PI homing vector (ZV). As before, ants were released at the centre of the test field for their first trial. Once the ants reach the rim (3.0 m radius around the release point) they were captured and attributed to one out of two visual conditions (U or N; Figure 1A). They were released either for a few seconds in U or N before being released again on the TF. Here again for the test, ants were captured and released on the test field for their second trial as ZV ants. For the unfamiliar transition where the centre of the test field is the ZV point, we forced the ant to run in the opposite direction of their exit point in the TF. This permits to keep the release point in the test field as the ZV point.

#### Recall experiment

During the final test of this experiment, ants released on the test field were no longer captured when reaching a 3.0 m distance from the release point. The search area was constrained with white barriers about 3 cm height, forming a square shape (6×6.0 m). For this experiment, we used ZV ants captured while exiting their nest and provided with a cookie to trigger homing motivation. ZV ants were released at the centre of the test field and free to explore. If during their exploration ants depart from the centre about 1.8 meters and return to the released point, they were captured again (at the centre of the test field). From there we created two groups. Group 1 ants were released at a novel location at least 1.8 m away from the centre. Group 2 ants were released again at the centre of the test field (the point of capture) (Figure 3C). After 4 repetitions of this procedure, ants were tested by letting them free to explore for at least 3 min.

### Data extraction Analysis of the ants’ direction of movement

Data was extracted and analysed have been run using the free software R (v 3.6.2. R Core Development Team). The recorded paths were digitized as (X, Y) coordinates using ImageJ. After digitization, all paths were discretized with a constant step length of 0.02 m. Descriptive statistics of this path such as the mean direction of movement (mu) as well as the mean circular vector length (r, a measure of dispersion) were extracted on R with the package Circular. The mean directions (mu) were analysed using a Rayleigh test (from R package: Circular) that also includes a theoretical direction (analogous to the V-test). To test whether the angular data are distributed uniformly as a null hypothesis or if they are oriented toward the theoretical direction of the nest as indicated by the state of the PI.

For the recall experiment, we did a density heat map of the ants’ position during the first trial and the test for group 1 and 2. The density maps have been done on the first 28 m of the test path. Path have been discretized into an X/Y matrix representing the space of the test field with a resolution of 25 cm.

We thus obtain with which frequency the ant has been in each square (0.25*0.25 cm^2^). From this matrix, we calculated an exploratory index (Fig; 3F) that correspond to the number of squares explored divided by the distance walked (28 m). This index between groups has been compared by using a one-tails permutation test.

Then we computed a density map at the population scale (Fig.3 D) by summing individuals density map within each group. To facilitate meaningful comparisons between the two groups, we normalized the heat maps accordingly.

### Data Time Warping (DTW) analysis

To reveal the occurrence of identical pattern between paths, we run a Dynamic Time Warping (DTW) analysis (from R package: SimilarityMeasures). This analysis allows for a comparison of two X, Y trajectories. The DTW computes a similarity value that corresponds to the sum of the distance between each point of trajectories. High values indicate that the two trajectories are rather dissimilar, while low values indicate that both trajectories are similar.

Rather than comparing the entire path, we compared segments between two trajectories of the same ants. To extract a segment, we choose a focal path. Then we indexed a focal point p0 and extracted the corresponding X, Y coordinates. These coordinates served to identify the closet point p1 in the subsequent path. We extracted a segment of 1.2 m around these focal points p0 and p1. We choose a focal point p0 every 10 steps. Between all segments, we did a DTW analysis. We used the median of all DTW within two paths for statistical tests.

To test whether the paths of ants were more similar within rather than between individuals, we compared the path of trial 1 of an individual to the path of trial 2 of all other ants. Then we obtain the median of these DTW analysis value across all ant comparisons. Medians of DTW values were then compared between the ‘between individuals’ and ‘within individuals’ paths comparisons using a permutation test for paired data.

### Computational model

We designed an agent-based model (Figure 4) using Python (Version 3.7.4). The agent consists of a point with position x,y and an associated heading orientation θ (we inferred that ants do not walk sideways and therefore heading θ is defined as the direction between the previous and current point location). The model runs in discrete time steps, with each time-step representing an iteration of the algorithm. The test field was modelled as a 3 m radius X,Y space in which the agent could update its position with a step length of 0.02 cm at each time step.

#### Agent memories

The agent’s memories of its routes on the test field were modelled as a 3D matrix, representing the space of the test field. Two dimensions correspond to the X,Y space of the field (steps of 10 cm) while the third dimension corresponds to all possible headings θ for each location (0°è360°, steps of 5°). At first, the 3D memory matrix is filled with 0 values (i.e., unfamiliar) representing the absence of memory for any location*heading. As the agent explores the world, at each time step the memory matrix values increase towards 1 (i.e. familiar) for the X,Y,θ (position*heading) of the agent, based on a defined learning rate (see table 1). To account for the fact that a view memorized at a given X,Y, θ can be recognized at neighbouring locations*headings^24,83,84^, the familiarity value was diffused around the X,Y, θ in the memory space following an exponential decay towards 0 (see ‘diffusion’ parameters in Table 1). Note that the value of the diffusion parameter in X,Y captures the visual openness of the environment: because more open environments visually change slower with displacement, a given learnt view can be recognized from further away. Two such memory matrixes were built and updated in parallel: one representing the persistent memory PM and on the transient memory TM.

**Table 1.** Default parameters used for both models. The first column corresponds to the parameter name. The second column to their description. The third and fourth column are the actual value used in different simulation.

| Parameters | Description | Continuous learning Experiment (fig. 4 B,C) | Rewind Experiment (fig. 4 D,E) |
| --- | --- | --- | --- |
| Learning rate | Familiarity value for the current position*orientation in memory matrixes are updated such as:<br>New value = Old value + (1 – Old value) *learning rate. | TM = 0.35<br>PM = 0.35 |  |
| Weight PI | Weight of direction indicated by the PI. | 0.015 |  |
| Weight view | Weight of the direction indicated by the recalled familiarity. | Range between 0 and 1 |  |
| Weight inertia | Weight of the tendency to continue straight | 0.8 |  |
| Diffusion accros location (X,Y) | Parameters of the exponential law for memory diffusion. | 0.6 Meaning that the memory diffuses 40 cm on either side of the position. | Open: 0. 5; spill = 50 cm.<br>Cluttered: 1; spill = 20 cm. |
| Diffusion across heading ( $\theta$ ) | Parameters of the exponential law for memory diffusion. | 0.7 Meaning that the memory diffusion spill around 25 degrees around the orientation. | |
| Step length | The distance made by the agent at each step | 0.02 meters |  |

#### Training phase (Trial 1)

For trial 1, the naïve agent (both TM and PM memory matrix initially set at 0) was moved along one of the paths of our real ants recorded during their first trails as ZV (N =23) while updating both its PM and TM memory matrixes.

#### Test phase (Trial 2)

We realized 10 replicas of the test trails. For each replica, the trained agent’s TM memory matrix was reset to 0, but the PM memory matrix values acquired during the training phase persisted. For One test (replicas), the agent was positioned at the middle of the test field in a random heading θ and let to guide its second path. At each time step, the agent performs a 0.02 cm step in a novel heading θ, which is determined by three directional forces integrated together as in vector-based navigation ^85,86^:

1. <u>Familiarity vector:</u> Because learnt view recognition is dependent on both position and orientation^24^ the agent extracts the ‘familiarity’ values of its current position and heading X,Y, θ in space by sampling the corresponding X,Y, θ values of both its PM and TM memories. Because TM inhibit PM, the ‘recalled familiarity’ is then obtained as ‘PM (X,Y, θ) – TM (X,Y, θ)’, and provides the weight (norm) of the familiarity vector for the current time step. To account from the fact that ants can learn view in multiple orientation at a same position and can associate these views to the most familiar route direction at this location^63^, the direction of the familiarity vector is obtained as the average of the θs familiarity of that location (X,Y) in the PM. To account for biological noise in retrieving the direction of a learnt route, a directional noise is added to this vector’s direction, drawn randomly from a Gaussian distribution of ± SD = 5°.
2. <u>Path Integration vector:</u> Path Integration (PI) continuously attracts homing ants towards their so-called Zero-Vector point location, usually at the nest position ^18^. Here we modelled ZV ants (captured at their nest) displaced to the centre of the test field, therefore, their PI vector always points from the agent’s current X,Y position to the centre of the test field. To account for biological noise in computing a PI home vector, a directional noise is added to this vector’s direction, drawn randomly from a Gaussian distribution of ± SD = 5°. The weight (norm) of this vector was fixed at 0.015 (see explanation below).
3. <u>Inertia vector:</u> To account for the fact that running ants are not constantly stopping and reassessing their orientation, we added an inertial vector, which direction correspond to the current heading θ of the agent. To account for biological motor noise, a directional noise is added to this vector’s direction, drawn randomly from a Gaussian distribution of ± SD = 10°. The weight (norm) of this vector was fixed at 0.8 (see explanation below).

At each time step, the new heading of the agent θ is obtained by summing the three vectors mentioned above (note that the norm of each vector function therefore as a weight, mimicking the weighted integration of directions achieved by ants^67,68,86,87^. The agent would then perform a step of 0.02 cm in this new θ to arrive at a new X,Y location in the test field, and both PM and TM memory matrixes would be updated to account from the ongoing learning of the current location.

Note that the familiarity weight of the current view varies from 0 (unfamiliar) to 1 (perfectly familiar). The weight of the PI and Inertial vectors have been manually selected to ensure that the agent did not follow routes more consistently than our real ant data (Figure S4B, C). Note that an agent guided only by the PI and inertia vectors (as in an exploration of a novel environment) would spontaneously display a systematic search centred on the PI Zero Vector point, as observed in ants ^33–36^.

### Modelled agent path analysis

The path model was analysed using R (version 3.6.2). To access to the similarity between two successive trajectories, we conducted a DTW (see method: “DTW analysis”). The median of the DTW between the first (training) and second trial (test) was extracted, as for the real ant data.

We compared DTW obtained from our ant’s path with the ones obtained from ants agents’ path (Fig. S4) using linear mixed models with the ant identity as random variable. It is important to note that as we realized 10 replicas of the test path (see *Test phase)* for the agent, we average the DTW value through replicas.

To estimate if ant agent indeed stays stuck in a particular trajectory pattern, we used a focal segment of the training path (trials one) with multiple other segments in subsequent test paths (trials two) to determine if the agent has followed a similar pattern. The comparison is made if the centre-to-centre distance between two segments is less than 60 cm and has a similar direction (within 25 degrees). The number of resulting comparisons is then used as a proxy for the number of times the agent repeated the pattern of the trial’s one.

#### Rewind experiment

To determine if our model could replicate behavioural signatures observed in the literature, we virtually reproduced an experimental paradigm where ants are forced to repeat a route segment several times consecutively. Typically, repetition after repetition, ants typically stop following their route and start to meandering away ^69,70^.

We replicated this experimental paradigm. First, we conducted 50 training sessions where the ant agent was trained along a direct route from the feeder to their nest located 7.0 m away. During training, the agents navigate with a home vector and progressively update their memory banks.

For a first test, we aimed to investigate the specific effect of visual repetition by re-running one portion of the route. We forced the agent to repeatedly traverse the same portion of the route by systematically catching the agent when reaching 1 m from its virtual nest and releasing it at the beginning of the inbound trip again. Between each repetition, as the agent does not encounter novel views, the TM is not reset, and the agent therefore accumulates familiarity in its TM memory across repetition. We did 6 replicates of this procedure for 20 ants’ agents. We then analysed paths, by using a sinuosity index (The inverse of the mean resultant vector calculated over the entire path). We then compared this value between the first and the last trials with a Wilcoxon test for paired data.

Lastly, we explored how the environmental structure influences the impact of route repetition on route disturbance. To achieve this, we modified the memory diffusion along the X and Y axes (see Table 1). Because more open environments visually change slower with displacement, a given learnt view can be recognized from further away, which can be modelled by having the familiarity values diffusing further along the X,Y of the memory matrixes. We thus simulated cluttered areas by reducing memory diffusion, simulating scenarios where the ant agent moves through a cluttered scene that therefore changes rapidly with displacement (see Table 1). Note that because ants were tested as ‘zero-vector ant’ captured at their nest and released at the beginning of the route, their PI home vector points in a direction opposite to the route familiarity direction from the first trial onwards ^88^, to account for his effect the PI vector weight was fixed at 0.015.

## Acknowledgments

We express our gratitude to Xim Cerda and the helpful team at Spanish National Research Council (CSIC Seville) and appreciated the help during field work preparation and data collection of Joséphine Gouleau, and Christelle Gassama. We thank the ants tested in these experiments for their participation.

## Funding

European Research Council, Grant reference number: EMERG-ANT 759817, Author: Antoine Wystrach.

## Supplemental information

**Figure S1.**
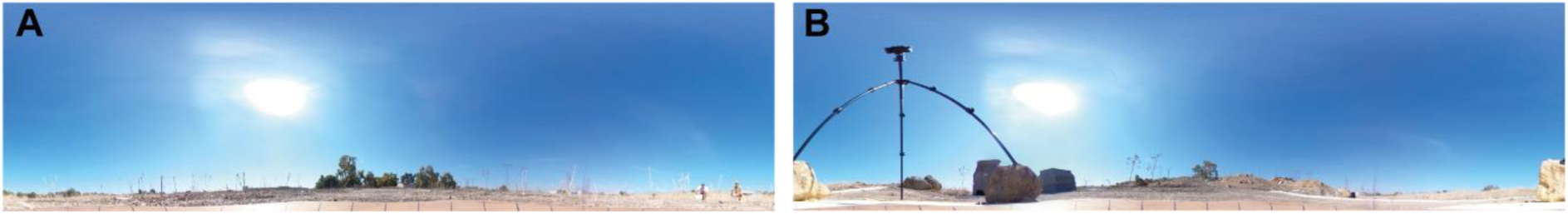
panoramic pictures. (A) The familiar route area and (B) the Unfamiliar test field 100 m away.

**Figure S2:**
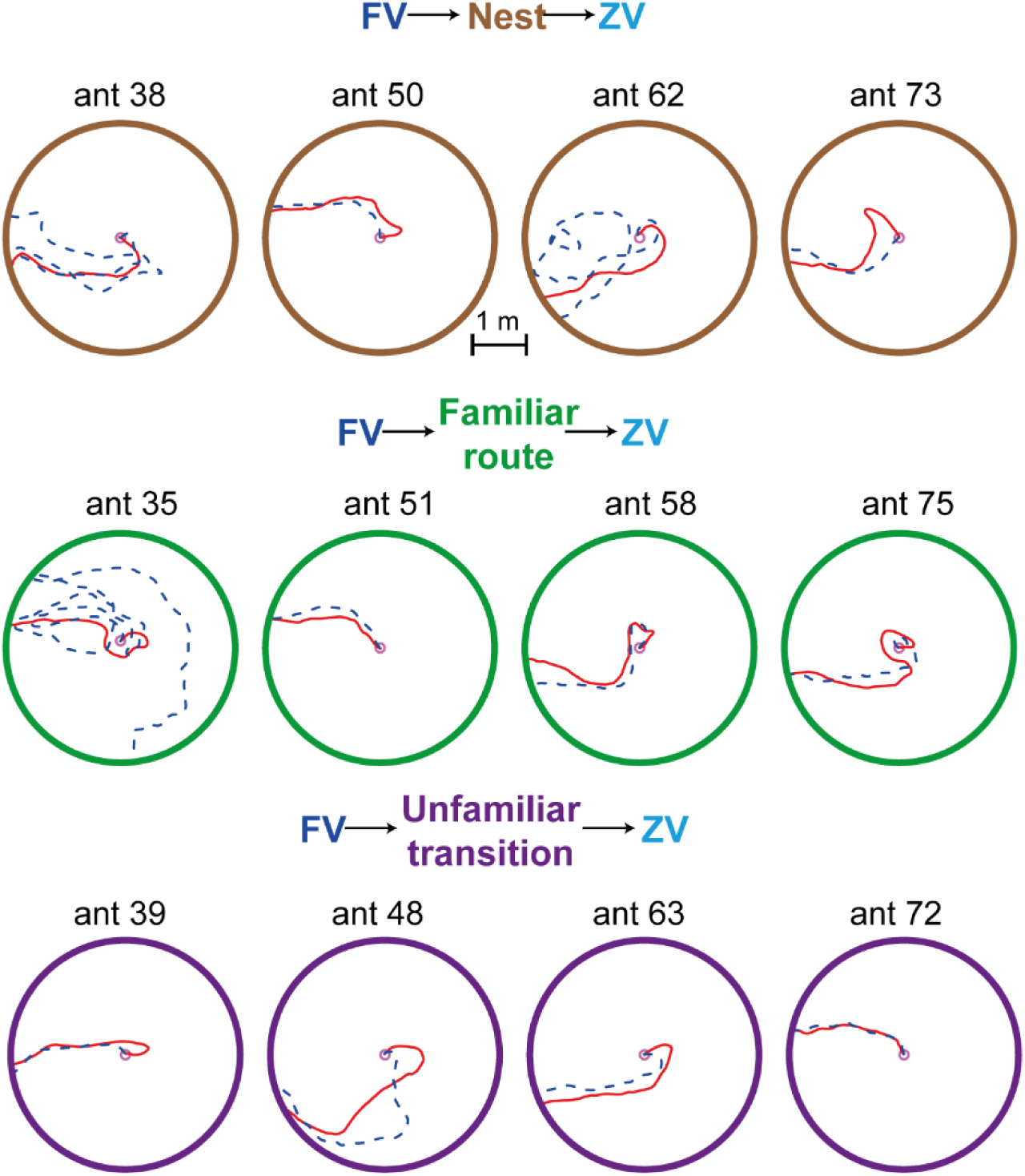
Experimental manipulations in FV Ants: samples Paths Path of ants trains as FV in the test field. After the first trials (red path), ants were either released at the nest between each recording session (first row), in their familiar route (second row) or in another unfamiliar place (third row). During the test – for all group-ants are recorded in ZV without interfering with their path.

**Figure S3:**
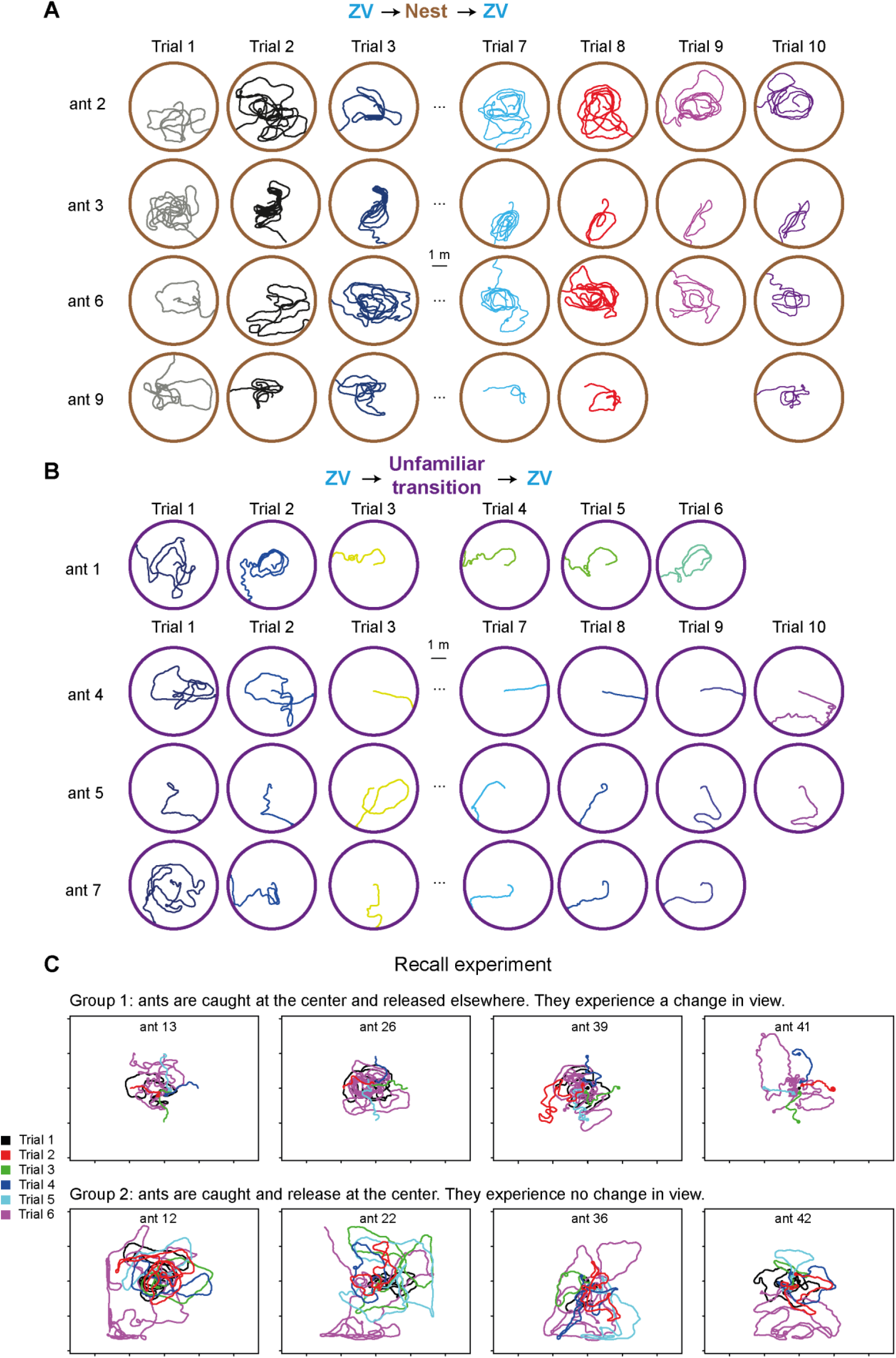
Experimental manipulations in ZV Ants: samples Paths. (A,B) Path of ants trains as ZV in the test field. After the first trials, ants were either released at the nest between each recording session (A) or in another unfamiliar place (B). (C) Example path of the “recall experiment”. For the first trial ZV ants are released and capture at the centre of the TF. Then for the next four trial ants are split into two groups that will experience either a changement of view (C, group 1) or not (C, group2). Finally, during the test (sixth trial, purple path) – for both group-ants are recorded without interfering with their path. On these examples’ path we can clearly see an attraction for the centre for the group 1 since trials 2. While group 2 tend to continue and widen their search.

**Figure S4.**
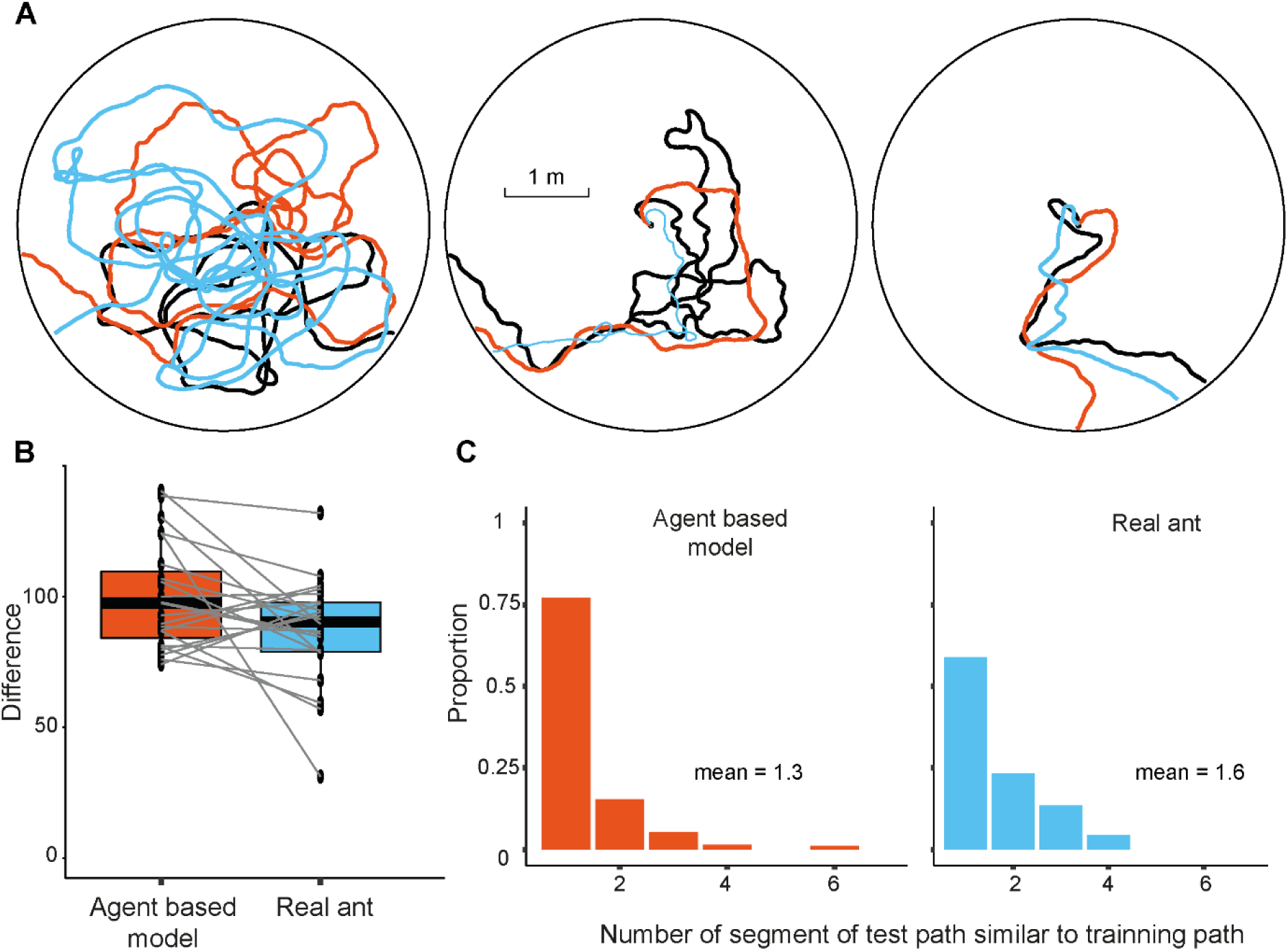
The feedforward inhibition model prevents to get stuck in a looping path signature. (A) Example path of the test path for model (orange) and real ant data train in ZV learning protocol (blue). The model has been trained once on the first path (see method) of real data (black path). (B) Comparison of the Similarity analysis (DTW) between models and real ants’ data trained as ZV (see method). Each path has been cut into segments of 2.4 meters, and each segment has been compared with the closest neighbour segment in a subsequent trial to estimate their similarity. Data obtained from the models has been average through replica. (C) Show the distribution of overlapping segments between the test path (trial two, A, coloured path) and training path (trial one, A, black path) for both model and real ant data. We compared one training path segment (trials one) with multiple test path segments (trials two). Two comparable segments are defined if their centre-to-centre distance is less than 60 cm and have a similar direction (within 25 degrees). The number of comparisons was used to estimate how many times a portion of route from trials one was repeated during the test path. We selected this range of parameters as the model crucially not follow route better than our real data (B, Anova < 0.05, post hoc analysis ps <0.05, mean DWT and SD: model = 99, 20; real ant data = 87, 20).

## Notes

### Competing Interest Statement

The authors have declared no competing interest.

